# Data variability in standardised cell culture experiments

**DOI:** 10.1101/2021.02.27.433153

**Authors:** Ian G. Reddin, Tim Fenton, Mark N. Wass, Martin Michaelis

**Affiliations:** School of Biosciences, University of Kent, Canterbury, UK

## Abstract

Despite much debate about a perceived ‘reproducibility crisis’ in the life sciences, it remains unclear what level of replicability is technically possible [1,2]. Here, we analysed the variation among drug response data of the NCI60 project, which for decades has tested anti-cancer agents in a 60-cell line panel following a standardised protocol [3]. In total, 2.8 million compound/cell line experiments are available in the NCI60 resource CellMiner [4]. The largest fold change between the lowest and highest GI50 (concentration that reduces cell viability by 50%) in a compound/cell line combination was 3.16 x 10^10^. All compound/cell line combinations with >100 experiments displayed maximum GI50 fold changes >5, 99.7% maximum fold changes >10, 87.3% maximum fold changes >100, and 70.5% maximum fold changes >1000. FDA-approved drugs and experimental agents displayed similar variation. The variability remained very high after removal of outliers and among experiments performed in the same month. Hence, our analysis shows that high variability is an intrinsic feature of experimentation in biological systems, even among highly standardised experiments in a world-leading research environment. Thus, a narrow focus on experiment standardisation does not ensure a high level of replicability on its own.

## Introduction

In the life sciences, there is a crisis narrative and a perception of a lack of data reproducibility (“reproducibility crisis” or “replication crisis”) [2,5–10]. However, the actual scale of the crisis remains unclear and evidence is largely anecdotal [2]. Much of the data are based on researcher views expressed in survey responses [11–13] or provided as Comments or Correspondence without providing detailed information [14,15].

One project that investigates the replicability of research data is the ‘Reproducibility Project: Cancer Biology’, which independently repeats influential preclinical studies (https://elifesciences.org/collections/9b1e83d1/reproducibility-project-cancer-biology). So far, 17 replication studies have been completed. Five studies reported the successful reproduction of the original studies [16–20], while eight reported a mixed outcome [21–28], and four failed to reproduce the original findings [29–32]. It is not clear whether these data are representative. It is a small dataset focused on small, early, and highly cited studies, which are more likely to overestimate effects [33].

There is also a lack of agreement on the expected level of data replicability [1]. In a dispute about the consistency of two large pharmacogenomic screens in cancer cell line panels, the Genomics of Drug Sensitivity in Cancer database and the Cancer Cell Line Encyclopedia [34–37], four analyses by different groups concluded a reasonable level of consistency [37–40], while six studies, all by the same group, disagreed [36,41–45].

To develop a realistic understanding of the replicability of standardised assays in a world-leading research environment, we investigated the variation in drug response data from the NCI60 screen [3]. Since 1985, the NCI60 screen has tested the anti-cancer activity of thousands of compounds multiple times in a 60-cell line panel following strict standard operating procedures (Figure 1A) [3,4,47–51]. Thus, the NCI60 database provides an unprecedented wealth of data on the replicability of findings using highly standardised procedures by highly skilled experts.

**Figure 1.**
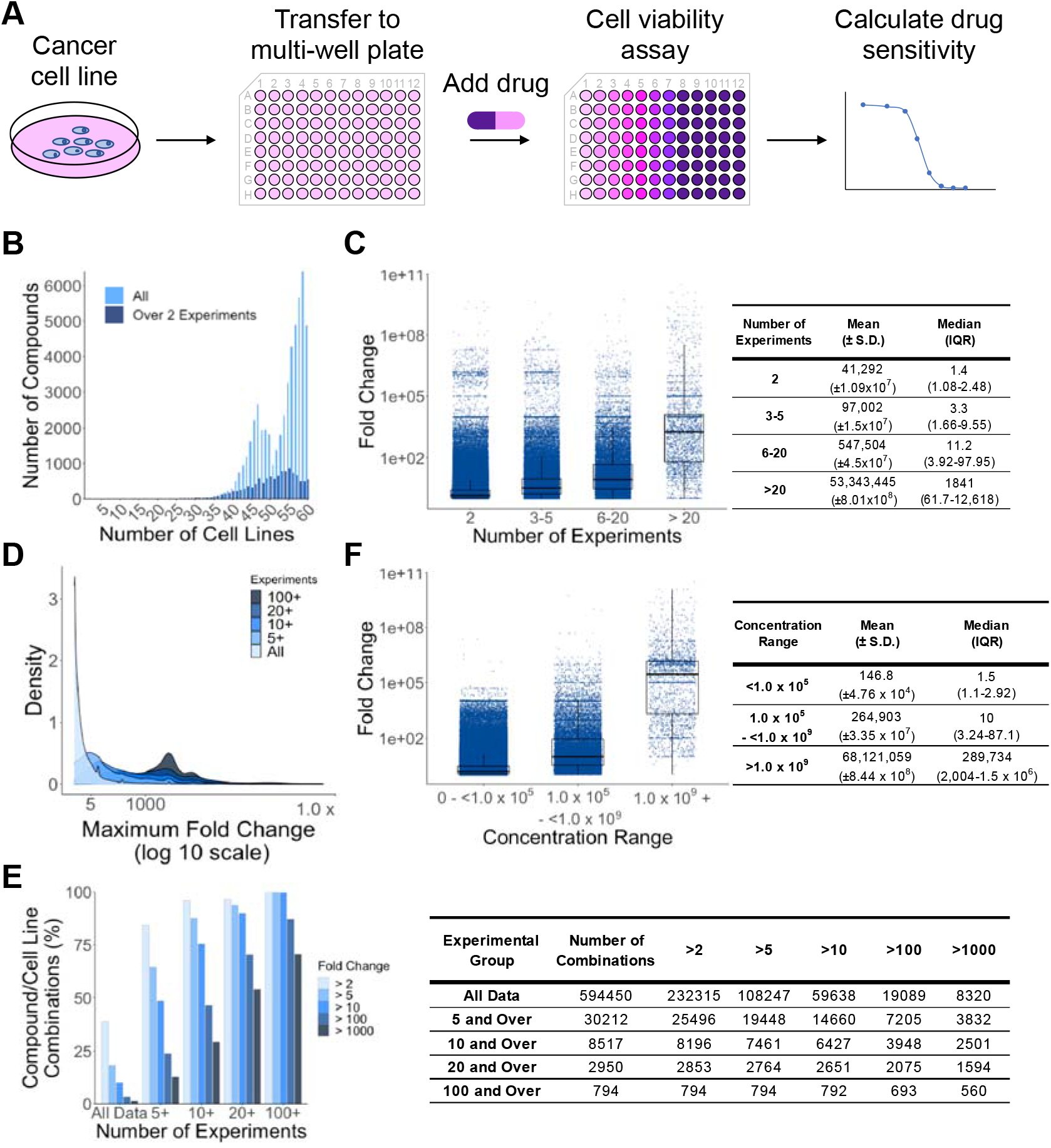
Variability in NCI60 GI50 data. **A)** Overview of the principle of the NCI 60 screen. **B)** Compound/cell line combinations with two or more experiments in the NCI60 database. **C)** GI50 fold changes in dependence on the number of experiments per compound/cell line combination. Numerical data are presented in the adjacent table. **D)** Distribution of maximum GI50 fold changes illustrated by density plots for experimental compound/cell line combination groups with an increasing minimum number of experiments. **E)** Percentage of compound cell line combinations with maximum fold changes above the indicated thresholds in dependence of the number of experiments. Numerical data are presented in the adjacent table. **F)** Distribution of GI50 fold changes in dependence of the concentration ranges in which compounds were tested. Numerical data are presented in the adjacent table.

## Results

### NCI60 drug response data are characterised by a high level of variability

All drug sensitivity data derived from NCI60 testing are made available via Cell Miner [4,47,48,50,51]. In total, 52,585 compounds were tested in the NCI60 resulting in 2.8 x 10^6^ compound/cell line combinations. Two or more (up to 2,286) experiments were carried out for 11,841 compounds and 594,450 compound/cell line combinations (Figure 1B, Extended Data Table 1; Extended Data Table 2). More than 100 experiments in at least one cell line were performed for 18 compounds and more than 1,000 experiments for two compounds (Extended Data Table 3). Concentration ranges varied from 10^1.2^ to 10^12.1^. 612 compounds were screened with multiple concentration ranges, and the most common concentration range was 10^4^ (11,213/ 94.7% of the compounds), representing the standard testing range using five 10-fold dilution steps (Extended Data Table 4).

The maximum fold change between the lowest and highest GI50 concentration (reduces cell viability by 50%) was detected for cyanomorpholinodoxorubicin in the colorectal cancer cell line COLO 205 (3.16 x 10^10^) (Extended Data Figure 1A, Extended Data Table 5). 232,315 (39.1%) drug/cell line combinations displayed maximum fold changes >2, 108,247 (18.2%) drug/cell line combinations fold changes >5, 59,638 (10%) drug/cell line combinations fold changes >10, 19,089 (3.2%) drug/cell line combinations >100, and 8320 (1.4%) drug/cell line combinations >1,000 (Extended Data Table 5).

### Variability increases with the number of experiments

The percentage of compound/cell line combinations with high maximum fold change strongly increased with the number of experiments (Figure 1C, Extended Data Figure 2A, Extended Data Figure 2B, Extended Data Table 6). The mean and median GI50 fold changes increased from 41,292 and 1.4 for compound/cell line combinations with two experiments to 53,343,445 and 1841 for compound/cell line combinations with >20 experiments (Figure 1C, Figure 1D, Extended Data Table 6).

When we considered compound/cell line combinations with a minimum of five experiments, 25,496 (84.4%) of 30,212 compound/cell line combinations displayed maximum fold changes >2 and 3,832 (12.7%) compound/cell line combinations >1000. For compound/ cell line combinations with >100 experiments, 100% of 794 compound/ cell line combinations displayed a maximum fold change > 5 and 70.5% (560 out of 794) displayed a maximum fold change >1000 (Figure 1E, Extended Data Table 7).

Taken together, maximum GI50 fold changes increase with the number of experiments. In agreement, a significant correlation was detected between maximum GI50 fold changes and the number of experiments per compound/cell line combination (Spearman correlation coefficient = 0.34, p < 2.2 x 10^−16^) (Extended Data Figure 3A).

### Variability increases with the concentration range covered

The observed fold changes also reflected the tested concentration ranges per compound/cell line combination in addition to the number of experiments, i.e. the broader the range of concentrations that were tested, the larger was the maximum fold change (Figure 1D, Extended Data Table 8). A positive correlation was observed between concentration range and maximum fold change for all compound data (Spearman correlation coefficient = 0.31, p < 2.2 x 10^−16^) (Extended Data Figure 3B).

The mean and median GI50 fold changes for compound/cell line combinations for which a maximum concentration range <1.0 x 10^5^ was covered were 146.8 and 1.5, which increased to 68,121,059 and 289, 734 for those with a concentration range of ≥1.0 x 10^9^ (Figure 1F, Extended Data Table 8).

### Variability in FDA Approved Drugs

Since reliable clinical therapy outcomes depend on reproducible drug effects, it may be speculated that FDA-approved drugs are more robust in their drug response data than experimental agents. However, the drug response data observed for FDA-approved drugs displayed a similar variability like that observed across all tested compounds.

The NCI60 database contained data on 181 FDA-approved drugs, which had been tested at least twice, resulting in 399,686 experiments investigating 9,970 individual drug/cell line combinations (Extended Data Table 1). The number of experiments for drug/cell line combinations ranged from 2 to 2,286.

The maximum GI50 fold change was 1.25 x 10^10^ observed for mithramycin in four cell lines, the colorectal cancer cell line COLO-205 (26 experiments) (Extended Data Figure 1B), the CNS cell lines SF-295 (24 experiments) and U251 (27 experiments), and the ovarian cancer cell line IGROV1 (26 experiments). Mithramycin, a member of the aureolic acid family was approved in 1970 but only temporarily used for testicular carcinoma and other types of cancer due to serious side effects [52]. The second highest GI50 fold change (7.28 x 10^7^) was detected for paclitaxel, a stabilising tubulin-binding agent and one of the most commonly used anti-cancer drugs [53], in MDA-MB-435 (Extended Data Figure 1C), which had originally been assumed to be a breast cancer cell line, but was later found to be derived from the melanoma cell line M14 [54].

The maximum GI50 fold changes were higher among the FDA approved drugs than for the non-FDA approved compounds (Figure 2A, Extended Data Table 9), probably because they were tested in more experiments and at bigger concentration ranges (Figure 2A).

**Figure 2.**
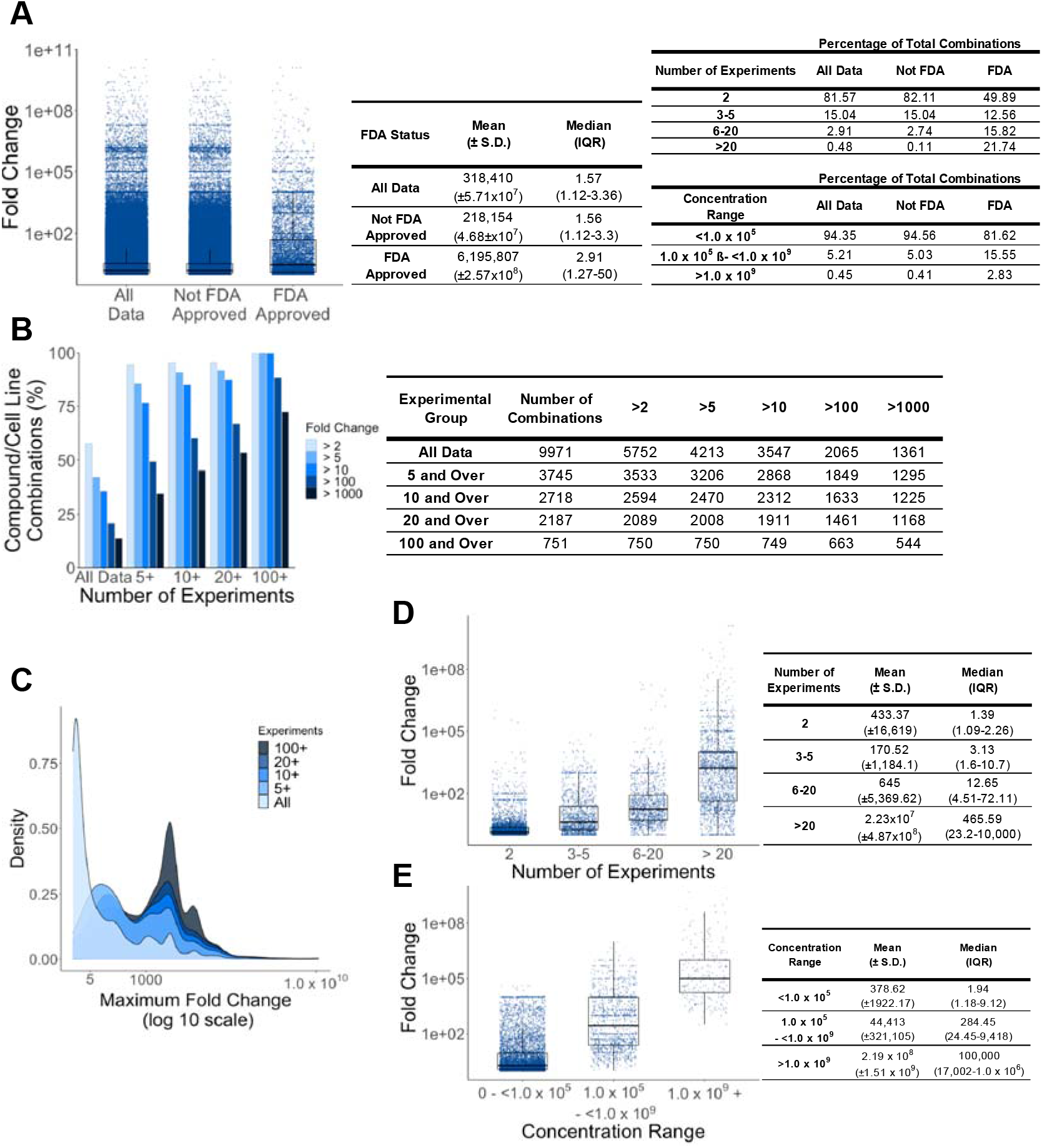
GI50 variation for FDA-approved drugs. **A)** Compound/cell line combinations with 2 or more experiments in the NCI60 database. Numerical data are presented in the adjacent tables. **B)** Percentage of FDA-approved drug/ cell line combinations with maximum fold changes above the indicated thresholds in dependence of the number of experiments. Numerical data are presented in the adjacent table. **C)** Distribution of maximum GI50 fold changes illustrated by density plots for experimental compound/cell line combination groups with increasing minimum numbers of experiments. **D)** GI50 fold changes in dependence on the number of experiments per compound/ cell line combination. Numerical data are presented in the adjacent table. (E) Distribution of GI50 fold changes in dependence of the concentration ranges in which compounds were tested. Numerical data are presented in the adjacent table.

When we considered the percentage of FDA-approved drug/cell line combinations with maximum GI50 fold changes >2, >5, >10, >100, and >1000 for combinations with >5, >10, >20, and >100 experiments (Figure 2B, Figure 2C, Extended Data Table 7, Extended Data Table 10), we obtained similar results to those across all compounds (Figure 1E).

In agreement with the findings across all compound/cell line combinations, the maximum GI50 fold changes also increased with experiment number when the FDA approved drug/cell line combinations were grouped into combinations with two experiments, 3 to 5 experiments, 6 to 20 experiments, and >20 experiments (Figure 2D, Suppl. Table 11), and the maximum GI50 fold change was also correlated with the number of experiments performed (Spearman’s correlation coefficient = 0.72, p < 2.2 x 10^−16^)(Extended Data Figure 3C).

Moreover, the maximum GI50 fold change increased with the concentration range covered (Figure 2E, Extended Data Table 12), and there was a significant correlation between the concentration range and the maximum GI50 fold change (Spearman’s correlation coefficient = 0.62, p < 2.2 x 10^−16^) (Extended Data Figure 3D).

Taken together, there is no indication that FDA-approved drugs would display less variability than experimental compounds.

### GI50 variability in experiments performed by month

The reproducibility of results may be affected by parameters such as changes in the reagents, e.g. use of different lots or batches, different experimenters, and using cell lines at different passages [2,55–57]. Hence, experiments performed closely together may be expected to display greater similarity than experiments performed at more distant points in time during the decades of anti-cancer compound testing by the NCI60.

To investigate the effects of the time of testing on data variability, we compared experiments performed in the same month to control samples of the same size that were randomly selected across the whole testing period. For this analysis, we used the 18 FDA-approved drugs that were tested at least 100 times in at least one cell line over periods of 95 to 275 months (Figure 3A, Extended Data Table 13), resulting in 51,872 drug/cell line combinations and in total 321,709 experiments (Figure 3B, Suppl. Table 14).

**Figure 3.**
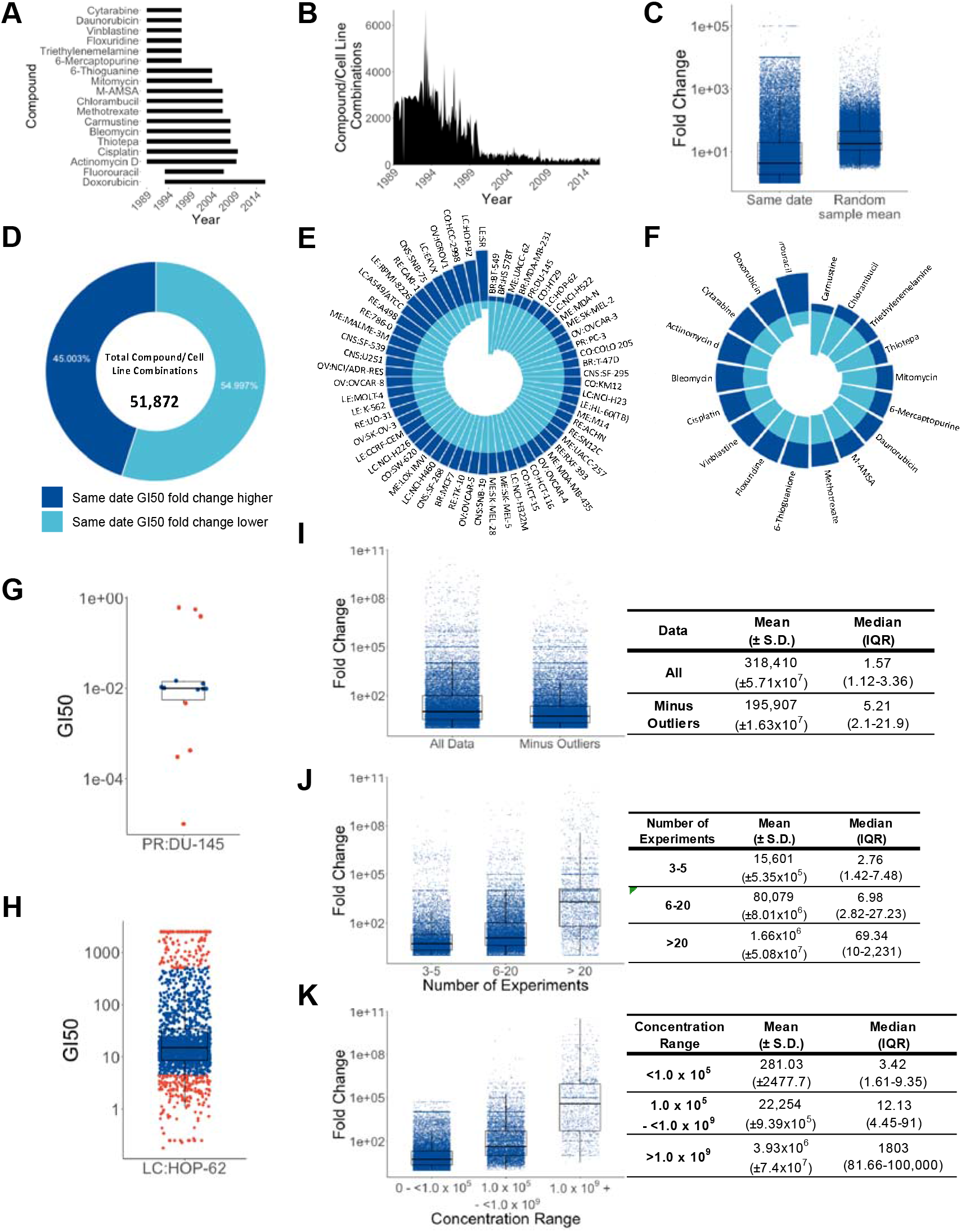
GI50 variability is high between compound/cell line combination experiments on the same date and is not caused by outliers. **A)** Time periods of drug testing for individual drugs. **B)** Testing of individual compound/cell line combinations by date. **C)** Maximum GI50 fold changes in experiments testing compound/ cell line combinations on the same date compared to maximum GI50 fold changes in 1000 random controls of the same sample size. **D)** Percentage of cases in which same date experiments had a higher fold change than control samples randomly picked across the timeline. **E)** Proportion of same date GI50 fold changes in compound/ cell line combinations that are higher or lower than random control samples per cell line. **F)** Proportion of same date GI50 fold changes in compound/ cell line combinations that are higher or lower than random control samples per drug. **G)** GI50 value distribution for maytansine in the prostate cancer cell line DU-145 (outliers indicated in red). **H)** GI50 value distribution for for **5-**fluorouracil in the lung cancer cell line HOP-62 (outliers indicated in red). **I)** Comparison of maximum GI50 fold changes before and after removal of outliers. Numerical values are presented in the adjacent table. **J)** Maximum GI50 fold changes increase with experiment number after removal of outliers. Numerical values are presented in the adjacent table. **K)** Maximum GI50 fold changes increase with the concentration range covered after removal of outliers. Numerical values are presented in the adjacent table.

For every set of experiments performed on the same date, we generated 1,000 random control samples of the same size and compared the value distribution. The variability of GI50 fold changes for same date experiments was indeed lower than for random control samples, but remained very high reaching up to 1.74 x 10^8^ (Figure 3C, Extended Data Table 15). Moreover, for 45% of the same date drug/ cell line combinations the GI50 fold change was higher than the mean fold change of the corresponding 1,000 random samples (Figure 3D, Extended Data Table 16).

When we looked at the data per cell line, the same date GI50 fold changes were higher than the mean random sample fold changes for the majority of drugs in ten cell lines, higher for half of the drugs in three cell lines, and lower for the majority drugs in the remaining 47 cell lines (Figure 3E, Extended Data Table 17). When we looked at the individual drugs, six displayed a majority of drug/cell line combinations with higher mean same date GI50 fold changes higher than in the random samples and twelve drugs displayed lower ones (Figure 3F, Extended Data Table 18).

Taken together, experiments performed in close timely proximity display lower variability than experiments performed over a longer time period, but the same date variability remains very high.

### High GI50 variability is not caused by outliers

Finally, we determined GI50 outliers for compound/cell line combinations with 5 or more experiments (738 compounds, 30,212 compound/cell line combinations, 598,243 GI50 values) using the adjusted boxplot method [58]. 5.7% (34,216) of GI50 values were outliers and 43.7% (13,208/30,212) of compound/cell line combinations had at least one GI50 outlier (Extended Data Table 19). This indicates that outliers are not responsible for the observed variability of GI50 in the majority (56.3%) of experiments.

The highest percentage of outliers was 50% (7/14 experiments for maytansine in DU-145 prostate cancer cells) (Figure 3G, Extended Data Table 19). The greatest number of outliers was 291 (16.8%) out of 1731 experiments for 5-fluorouracil in HOP-62 lung cancer cells (Figure 3H, Extended Data Table 19). Outlier number increased with the number of experiments for a compound/cell line combination with a Spearman correlation coefficient of 0.25 (p < 2.2 x 10^−16^) (Extended Data Figure 4). The removal of outliers reduced data variability, but the overall variability remained very high with a maximum GI50 fold range of 2.5 x 10^9^ detected for maytansine in the ovarian cancer cell line OVCAR-5 over 35 experiments (Figure 3I, Extended Data Table 19).

As detected in the analysis across all experiments, maximum GI50 fold changes increased with the number of experiments and the concentration ranges covered also after the removal of outliers (Figure 3J, Figure 3K, Extended Data Table 19, Extended Data Table 20). A significant correlation was observed between experiment number and maximum GI50 fold change with a Spearman correlation of 0.39 (p < 2.2 x 10^−16^) (Extended Data Figure 5A) and between concentration range and maximum GI50 fold change with a Spearman correlation of 0.47 (p < 2.2 x 10^−16^) (Extended Data Figure 5B).

### No drift in drug sensitivity over time

Cancer cell lines may display substantial changes in genotype and phenotype over time [55,59]. Hence, part of the variability observed in drug sensitivity may be the consequence of a shift in drug response over time. To investigate this, we established timelines of the GI50 values for the 18 compounds, which had been tested at least 100 times in one or more cell lines. This resulted in time lines for 1080 compound/cell line combinations with time frames ranging from 95 months (vinblastine, floxuridine, cytarabine, daunorubicin, 6-mercaptopurine) to 275 months (Doxorubicin) (Figure 3A, Extended Data Table 13).

Drug/ cell line combinations, in which the fold change between the mean GI50 on the first experimental date and the mean GI50 on the last experimental date was 50% or greater than the maximum GI50 fold change for the data points in between were considered as candidates for a drift in drug sensitivity. Only six (0.56%) out of 1080 drug/cell line combinations fulfilled these criteria (Figure 4, Extended Data Table 21).

The distribution of the individual GI50 values for three of the drug/ cell line combinations (floxuridine/ SK-OV-3, methotrexate/ BT-549, 6-mercaptopurine/ BT-549) did not indicate a GI50 shift over time (Figure 4A-C, Extended Data Table 21, Extended Data Table 22). For the other three drug/ cell line combinations (vinblastine/ T-47D, bleomycin/ K-562, M-AMSA/ MDA-MB-435) a drift in sensitivity appears unlikely but cannot be excluded based on the data (Figure 4D-E, Extended Data Table 21, Extended Data Table 22). However, such observations are very rare. Moreover, a phenotypic drift in a cell line would be expected to result in changes in sensitivity to more than one drug over time. Hence, the data provide no evidence suggesting that the drug sensitivity of individual cell lines may have changed over time. These findings may also reflect that the NCI60 uses cell lines within a window of 30 passages [60].

**Figure 4.**
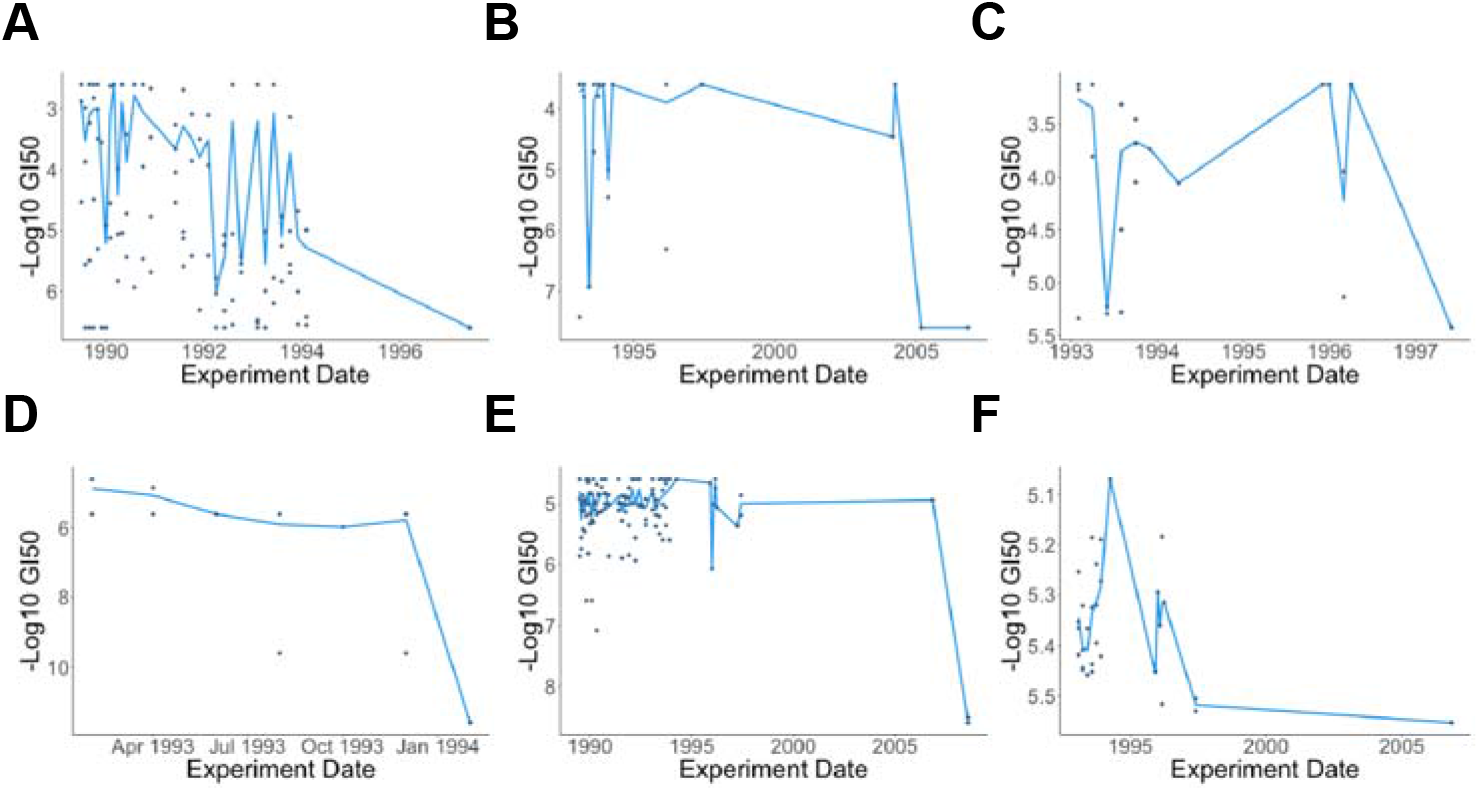
Experimental time lines for individual compound/cell line combinations. The six experimental timelines for compound/cell line combinations (with more than 100 experiments performed in at least one cell line) with a fold change between the first and last mean GI50 that is 50% greater than the maximum GI50 for the remaining experiments. **A)** floxuridine in the ovarian cancer cell line SK-OV-3, **B)** methotrexate in the breast cancer cell line BT-549, **C)** 6-mercaptopurine in the breast cancer cell line BT-549, **D)** bleomycin in the leukaemia cell line K-562, **E)** vinblastine in the breast cancer cell line T-47D, **F)** M-AMSA in the melanoma cell line MDA-MB-435. Plot lines represent mean GI50s while data points represent individual experiments at an experimental date.

## Discussion

To gain real-life insights into the extent of data variability in a standardised research environment, we here investigated the variation of GI50 values indicating drug sensitivity for compounds that had been tested multiple times in the NCI60 panel. The variation was large with the highest fold change between the lowest and highest GI50 in a given compound/cell line combination being 3.16 x 10^10^. As might have been expected, the fold change between the lowest and the highest GI50 in a specific compound cell line combination increased with the number of experiments and the concentration range tested.

CellMiner contains data on experimental compounds as well as on FDA-approved drugs that are in clinical use [4,46–51]. Although FDA-approved drugs might have been expected to result in more robust data, this was not the case and they displayed a similar data variability as that determined across all compounds. The variability also remained very high when we only considered experiments that were performed in the same months or removed outliers.

Given that the NCI60 uses highly standardised methods, a high variability of assay results appears to be an intrinsic feature of lab experiments. The large range of GI50 values observed for anti-cancer drugs is of potential relevance, given that cytotoxic anti-cancer drugs are typically used at maximum tolerated doses that cannot be further increased without unacceptable toxicity [61–63]. Moreover, the maximum effects of targeted drugs, e.g. antibodies or kinase inhibitors that interfere with cancer-specific structures or entities, do not further increase beyond the ‘optimal biological dose’, i.e. the dose at which the biological target is completely inhibited [61–64]. Hence, even a two-fold difference in the GI50, which occurred in 25,496 (84.4%) of 30,212 compound/cell line combinations with at least five experiments, is of potential relevance, as a two-fold increase of the clinical dose of an anti-cancer drug is rarely feasible.

Notably, there is awareness of this variability within the NCI60 project as indicated by strict quality control procedures in the presentation of NCI60 GI50 values in CellMiner, which result in the exclusion of up to 96% of experiments (48 out of 50) for a given compound/ cell line combination (Extended Data Table 23) [4].

However, such knowledge has not penetrated into scientific discourse. Many authors promote strict standardisation of experimental procedures as a strategy to improve data quality and reliability [2,5,15,45,65–71].

Our analysis of NCI60 data, which is of unprecedented depth and provided by a world-leading institution with unprecedented transparency, indicates that data variation remains very high even under ideal conditions that the vast majority of research groups will not be able to afford. Hence, our data suggest that experiment heterogenization, the testing of a hypothesis in many different (experimental) systems and datasets and different laboratories [72–79], is a much better strategy to generate robust and meaningful data.

In conclusion, our analysis demonstrates that the variation of experimental data is extremely high even under optimal conditions in a world-leading environment applying the highest standards. This shows that increased standardisation is not a straightforward way to resolve issues associated with limited replicability. Hence, increased data robustness will have to include additional strategies such as independent replication and experiment heterogenization, i.e. multiple testing of the same hypothesis using different approaches and models. Awareness of the inherent variability of experimental results, will help researchers to develop a realistic understanding of the meaning of their data and to design more diverse research strategies that will result in higher data robustness and reliability.

## Methods

### Data Availability

All data were obtained from CellMiner [80] Version 2.2. Dose concentration range data (June 2018 release) were obtained from the National Cancer Institute DTP NCI bulk data for download pages (https://wiki.nci.nih.gov/display/NCIDTPdata/NCI-60+Growth+Inhibition+Data). Of the 52,585 NCI codes given to compounds tested on the NCI-60 cell line panel, 42,794 were given to unnamed compounds and 9,791 were given to 9,027 named compounds. 262 named compounds received two or more individual codes. Some GI50 values represent minimum or maximum drug concentrations where the actual GI50 was not reached [4]. Since such values understate the actual data variation, we did not remove these data. All data generated during this study are included in this published article and its supplementary information files.

### Maximum GI50 fold change calculation

The drug sensitivity data was converted from −log10 GI50, to the GI50 (μM) for all compound, all cell lines and all experiments. Maximum fold changes were calculated for each compound/cell line combination with more than one experiment (594,450) by dividing the maximum GI50 for a cell line by the minimum GI50.

### Number of experiments and experimental groups

The number of experiments for each individual compound/cell line combination was calculated by counting all experiments performed on the same experimental date as well as experiments on different dates. The relationship between number of experiments and maximum fold change was investigated by using Spearman’s correlation coefficient as the distribution of maximum GI50 fold change was not normal.

The compound/cell line combinations were then assigned experimental groupings base on the number of experiments performed: all data, 5 or more experiments, 10 or more experiments, 20 or more experiments, and 100 or more experiments. This allowed comparison of “high” maximum fold changes (>2, >5, >10, >100, and >1000) for combinations with varied number of experiments. Additionally, compound/cell line combinations were assigned to experimental groups: 2 experiments, 3 to 5 experiments, 6 to 20 experiments and over 20 experiments. These experimental groupings enabled comparison of GI50 fold change statistics (mean, median, minimum, maximum, variance) for compound/cell line combinations with number of experiments ranging from lower to higher.

### Concentration range and experimental groups

Maximum dose concentration range for a compound/cell line combination was determined by using the minimum and maximum dose concentration used in an experiment for an individual compound on an individual cell line. The minimum concentration range was 1.0 x 10^1.2^ and the maximum concentration was 1.0 x 10^12.1^. The relationship between dose concentration range and maximum fold change was investigated by using Spearman’s correlation coefficient as the distribution of maximum GI50 fold change was not normal.

Compound/cell line combinations were assigned to groups based on the dose concentration range for that combinations: maximum concentration range less than 1.0 x 10^5^, maximum concentration range 1.0 x 10^5^ to 1.0 x 10^9^ exclusive and maximum concentration range 1.0 x 10^9^ and above. These experimental groupings enabled comparison of GI50 fold change statistics (mean, median, minimum, maximum, variance) for compound/cell line combinations between lower and higher concentration ranges.

### FDA-approved compound analysis

All compounds that were classed as FDA-approved drugs by the NCI-60 in CellMiner Database Version 2.2 and where two or more experiments had been performed were extracted from the complete dataset. This created an FDA-approved dataset of 181 drugs for which 399,686 experiments for 9,970 individual drug/cell line combinations were performed. Analysis of relationship between the number of experiments/concentration ranges and maximum GI50 fold change for drug/cell line combinations was performed as for the complete dataset, described above.

### Experiments on the same date

Month and year of each experiment was available so experimental timelines were established for compounds by calculating the time between the first and last experiment date. Multiple experiments were carried out on the same date for many of the compound/cell line combinations, particularly the 18 compounds with at least one cell line with 100 total experiments. The data for these 18 compounds, 17 of which were FDA-approved, was extracted from the complete dataset to create a subset of data deemed suitable to compare GI50 variability on the same date with GI50 variability over an experimental timeline.

The maximum GI50 fold change on each date where there were multiple experiments for a compound/cell line combination were calculated by dividing maximum GI50 by minimum GI50 value. The number of experiments on a specific date for a compound/cell line combination was used to determine the maximum GI50 fold change over the same number of experiments picked randomly from that combination’s experimental timeline. This was performed 1000 times so that for every compound/cell line combination and experimental date with a maximum GI50 fold change over multiple experiments there were 1000 corresponding maximum GI50 fold changes calculated from random samples of the same number of experiments on that compound/cell line combination’s timeline. The mean maximum GI50 fold change was calculated for the 1000 random samples for each compound/cell line combination and the number of maximum GI50 fold changes for experiments on the same date higher and lower than the random sample mean were counted. For each compound, significance of the difference between same date maximum GI50 fold change and sample mean GI50 fold change was calculated using Wilcoxon Rank Sum Test. This was performed using all cell line data combined for each compound and for each cell line individually for each compound. Where a significant difference between same date and random sample mean maximum GI50 fold changes were observed, the number of times the same date GI50 fold change was higher or lower than the random sample mean maximum GI50 fold change was counted.

### Drift in drug sensitivity

The mean GI50 fold change was calculated for each experimental date (month) for the 18 compounds with 100 or more experiments for at least one cell line. The GI50 fold change between the first experimental date and the last experimental date was calculated using the mean GI50 on those dates. The first/last GI50 fold change was then compared to the maximum GI50 fold change for each compound/cell line combination and considered a candidate for a drift in sensitivity if it was 50% or more of the maximum fold change.

### Removal of outliers

The adjusted boxplot method was used to identify outlier thresholds. This method was chosen as the data set was highly skewed. To use this method the medcouple *(MC)*, a robust measure of skewness, had to be calculated (where *X_n_ = {x_1_, x_2_, … , x_n_ }* represents data for every compound/cell line combination):

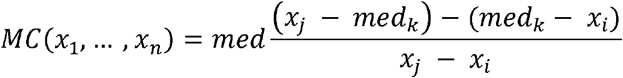

Where *med_k_* is the median of *X_n_*, and *i* and *j* have to satisfy *x_i_ ≤ med_k_ ≤ x_j_*, and *x_i_ ≠ X_j_*.

Using the *MC* the upper *(U)* and lower *(L)* thresholds could be determined. If *MC ≥ 0*:

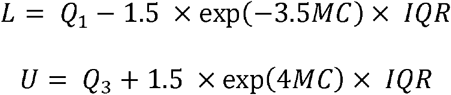

*MC ≤ 0*:

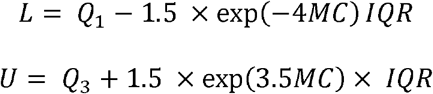

If *MC = 0* the adjusted boxplot method was not used but instead the Tukey method was used:

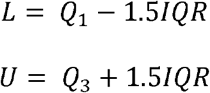

Where *Q_1_* is the lower quartile, *Q_3_* is the upper quartile and *IQR* is the interquartile range.

For each compound/cell line combination any GI50 value below *L* or above *U* were removed from the dataset. Analyses were performed on this dataset as previously described for the complete dataset.

### Data processing

Data was carried out using perl version 5.26.0, Microsoft Excel (2011) and R statistical packages version 3.4.4. Perl modules Statistics:Descriptive and Statistics::R were used. Packages used in R were robustbase, dplyr, webr, moonBook, tidyverse, reshape2, scales, gplots, ggpubr, ggExtra, RColorBrewer, corrplot, ggplot2, and tidyr.

## Author contributions

Data acquisition and analysis (I.R.), data interpretation (all authors), study conception (M.N.W., M.M.), study design (I.R., M.N.W., M.M.), manuscript drafting (I.R., M.M.), manuscript revision (all authors).

All authors have approved the submitted version and agreed both to be personally accountable for their own contributions and to ensure that questions related to the accuracy or integrity of any part of the work, even ones in which they were not personally involved, are appropriately investigated, resolved, and the resolution documented in the literature.

## Data availability

All raw data are available from CellMiner [80] Version 2.2. All other data are provided in the manuscript and its supplements.

