## Supplemental Figures for "Data variability in standardised cell culture experiments"

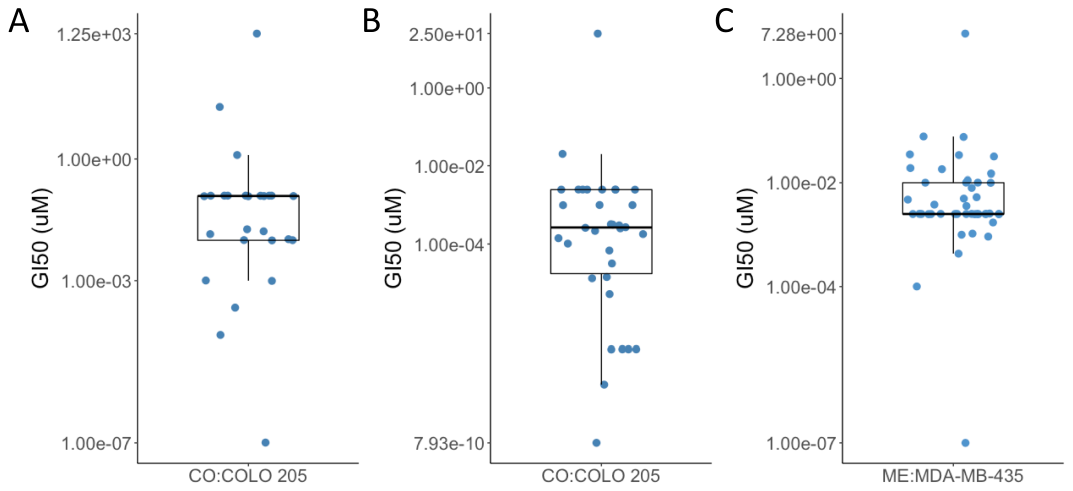

**Extended Data Figure 1. Individual GI50 values for compound/cell line combinations. A)** Cyanomorpholinodoxorubicin in the colorectal cancer cell line COLO 205. **B**) Mithramycin in COLO 205. **C)** Paclitaxel in the melanoma cell line MDA-MB-435.

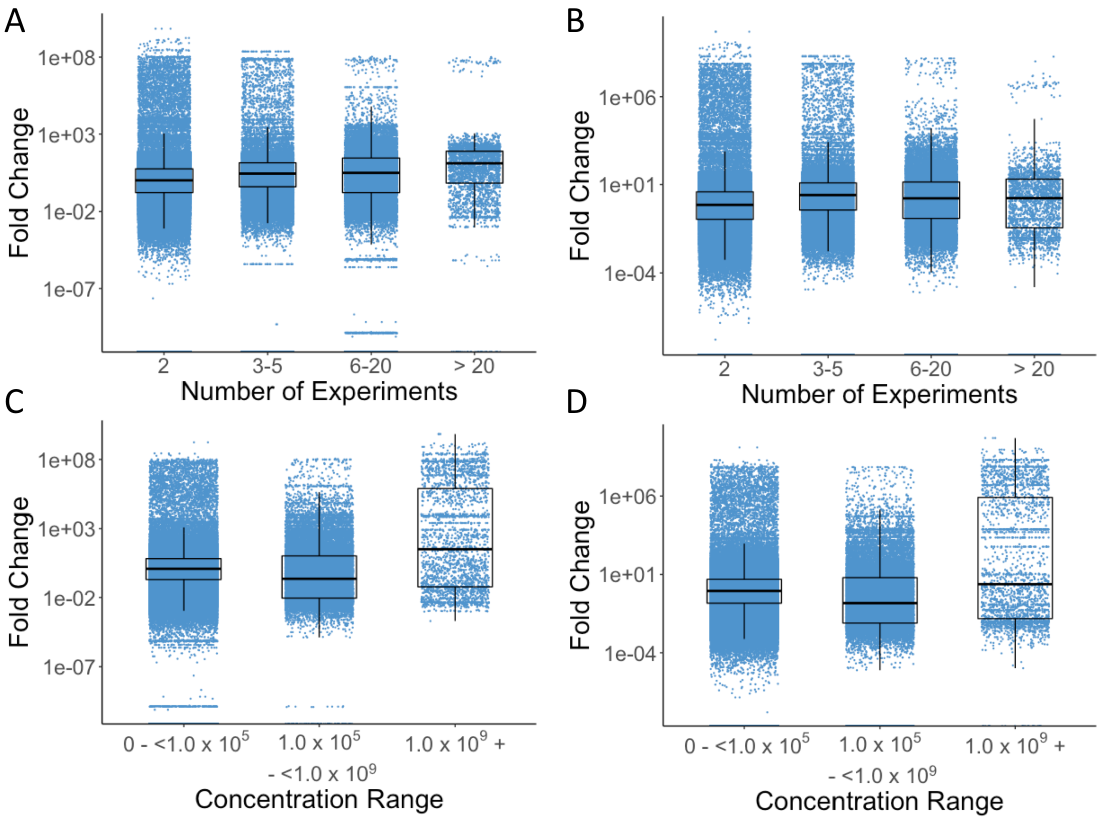

**Extended Data Figure 2. Standard deviation and interquartile range of GI50 values in different experimental and dose concentration range groups.** Standard deviation and interquartile range of compound/cell line combinations in increasing experimental groups **(A & B)** and increasing dose concentration ranges **(C & D)**.

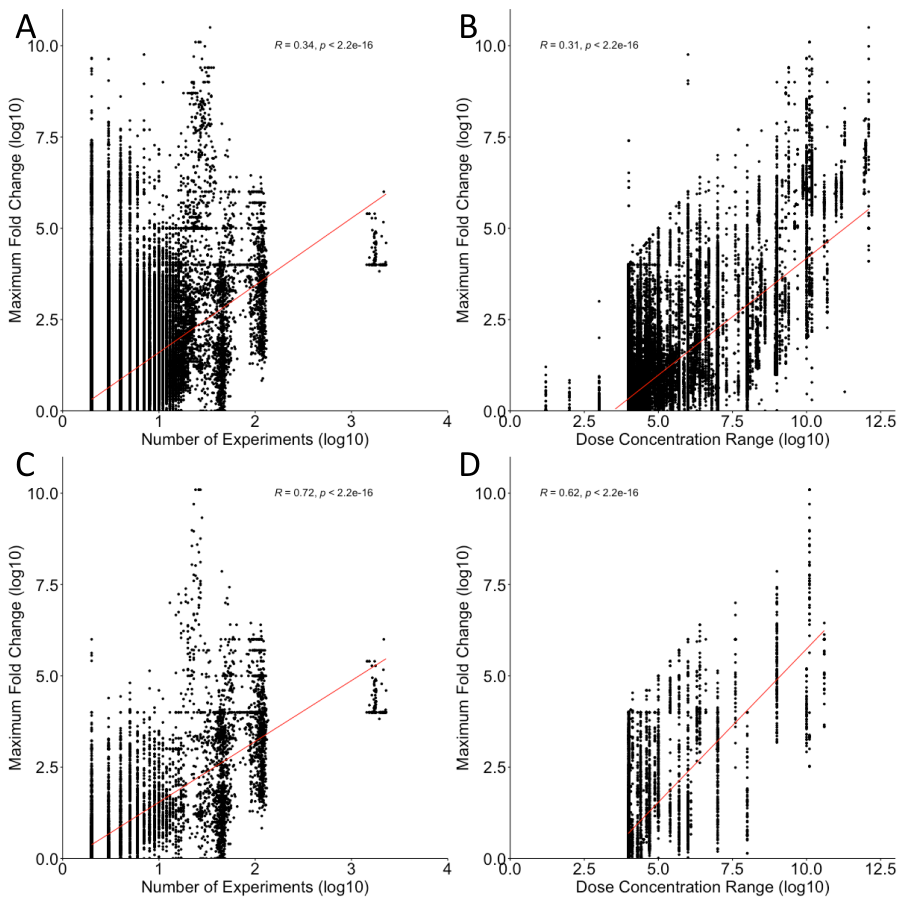

**Extended Data Figure 3. Variability increases with number of experiments and concentration range covered. A)** Spearman’s correlation between the maximum GI50 fold change for all compound/cell line combinations and the number of experiments. **B)** Spearman’s correlation between the maximum GI50 fold change for all compound/cell line combinations and the concentration range covered. **C)** Spearman’s correlation between the maximum GI50 fold change for FDA-approved drug/cell line combinations and the number of experiments. **D)** Spearman’s correlation between the maximum GI50 fold change for FDA-approved drug/cell line combinations and the concentration range covered.

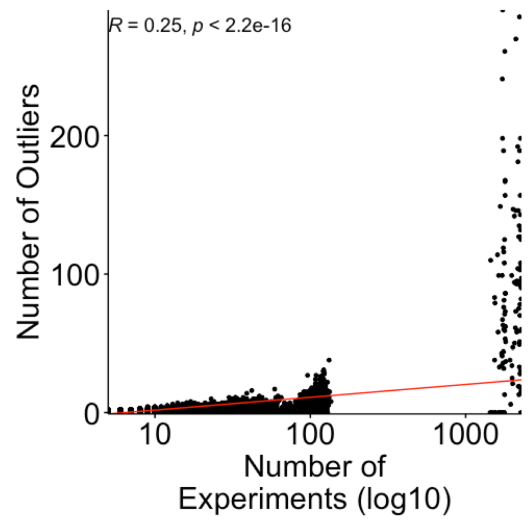

**Supplementary Figure 4. Number of outliers increases with the number of experiments.**

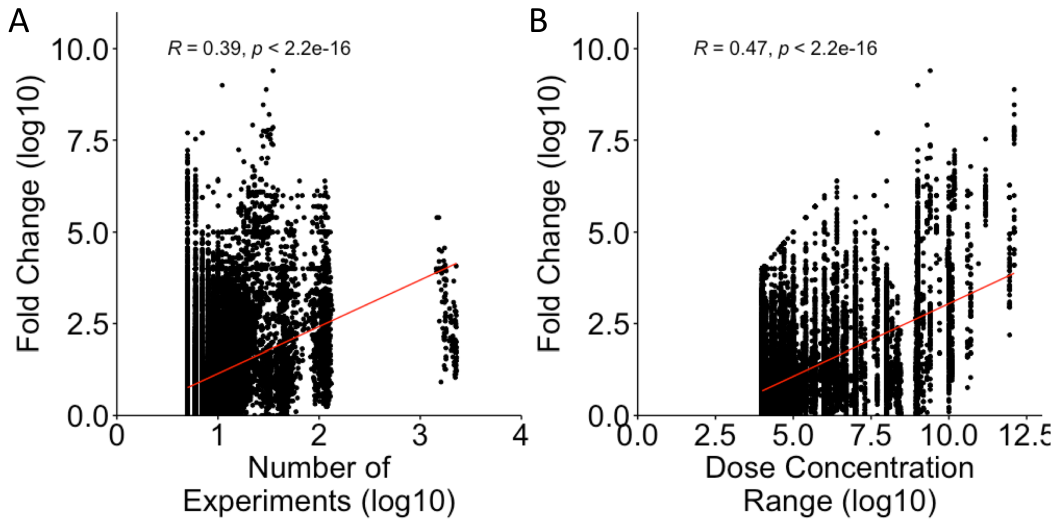

**Extended Data Figure 5. Variability increases with number of experiments and concentration range after removal of outliers. A)** Spearman’s correlation between the maximum GI50 fold change for compound/cell line combinations and the number of experiments. **B)** Spearman’s correlation between the maximum GI50 fold change for compound/cell line combinations and the covered dose concentration range.
